# An mTurquoise2-based glucose biosensor

**DOI:** 10.1101/2024.11.29.626064

**Authors:** Dennis Botman, Annemoon Tielman, Joachim Goedhart, Bas Teusink

**Author notes:** To whom correspondence may be addressed: Dennis Botman.

## Abstract

Glucose is an important substrate for organisms to acquire energy needed for cellular growth. Despite the importance of this metabolite, single-cell information at a fast time-scale about the dynamics of intracellular glucose levels is difficult to obtain as the current available sensors have drawbacks in terms of pH sensitivity or unmatched glucose affinity. To address this, we developed a convenient method to create and screen biosensor libraries using yeast as workhorse. This resulted in TINGL (Turquoise INdicator for GLucose), a robust and specific biosensor with an affinity that is compatible with intracellular glucose detection. We calibrated the sensor in vivo through equilibration of internal and external glucose in a yeast mutant unable to phosphorylate glucose. Using this method, we measured dynamic glucose levels in budding yeast during transitions to glucose. We found that glucose concentrations reached levels up to 1 mM as previously determined biochemically. Furthermore, the sensor showed that intracellular glucose dynamics differ based on whether cells are glucose-repressed or not. Finally, the human codon-optimized version (THINGL, Turquoise Human INdicator for GLucose), also showed a robust response after glucose addition to starved human cells, showing the versatility of the sensors. We believe that this sensor can aid researchers interested in cellular carbohydrate metabolism.

## Introduction

Glucose is a major substrate used by many organisms to obtain energy and carbon needed for cellular growth^1,2^. This sugar is transported into cells and -for the vast majority of organisms- metabolized via glycolysis to generate ATP. *Saccharomyces cerevisiae* (budding yeast) transports glucose via a set of hexose transporters (HXTs) that operate as facilitated diffusion carriers, i.e. they rely on concentration gradients for net flux, similar as the human glucose transporters (GLUTs)^3,4^. It was shown that such transporters are very sensitive to product inhibition, and hence, the concentration of internal glucose has large implications on the glucose uptake rate and hence, glycolytic flux. Using an elaborate population-based biochemical assay involving double radio-active labeling, the intracellular glucose levels have been measured in yeast after glucose addition, and was estimated at approximately 0.7 to 1.5 mM ^5,6^. This is in the range of the high-affinity carriers and fitted with the corresponding glycolytic flux^7^. However, the existing method does not allow for dynamic measurements, let alone single-cell measurements at the fast timescales that metabolism is operating at.

Intracellular glucose levels can potentially be assessed using fluorescent biosensors. Various biosensors for glucose have been described. These sensors include the FLIPglu FRET and GIP sensors^8,9^, the single-GFP based Glifon sensors^10^, the cpGFP-based iGlucoSnFR sensors^11^, the mApple-based Red Glifons^12^ and the cpYFP based FGBP_1mM_ sensor^13^. Yet, these sensors suffer from pH sensitivity (among other issues) making them unsuitable to study intracellular glucose levels in dynamic contexts as glucose import and startup of glycolysis affects pH levels in a cell (approximately from 7.1-6.5)^14–17^. Recently, an mTq2-based glucose sensor was developed showing promising results regarding pH robustness^18^. However, the affinity of that sensor is very high (∼14 μM) and cellular characterization is lacking, questioning its use for live cell imaging. Thus, measuring the dynamics of intracellular glucose levels inside single living cells is still challenging and therefore the intracellular glucose levels in dynamically-changing conditions are still largely unresolved. To address this, we developed a robust mTq2-based biosensor using a simple but unique screening method. The developed sensor, named TINGL (Turquoise INdicator for GLucose) is a bright monomeric sensor that has a fast response rate to glucose, is pH robust, bright and monomeric and is therefore an excellent tool to study intracellular glucose levels at a single-cell level.

## Material and methods

### Yeast strains

All yeast strains used in this study are listed in table 1.

**Table 1.**
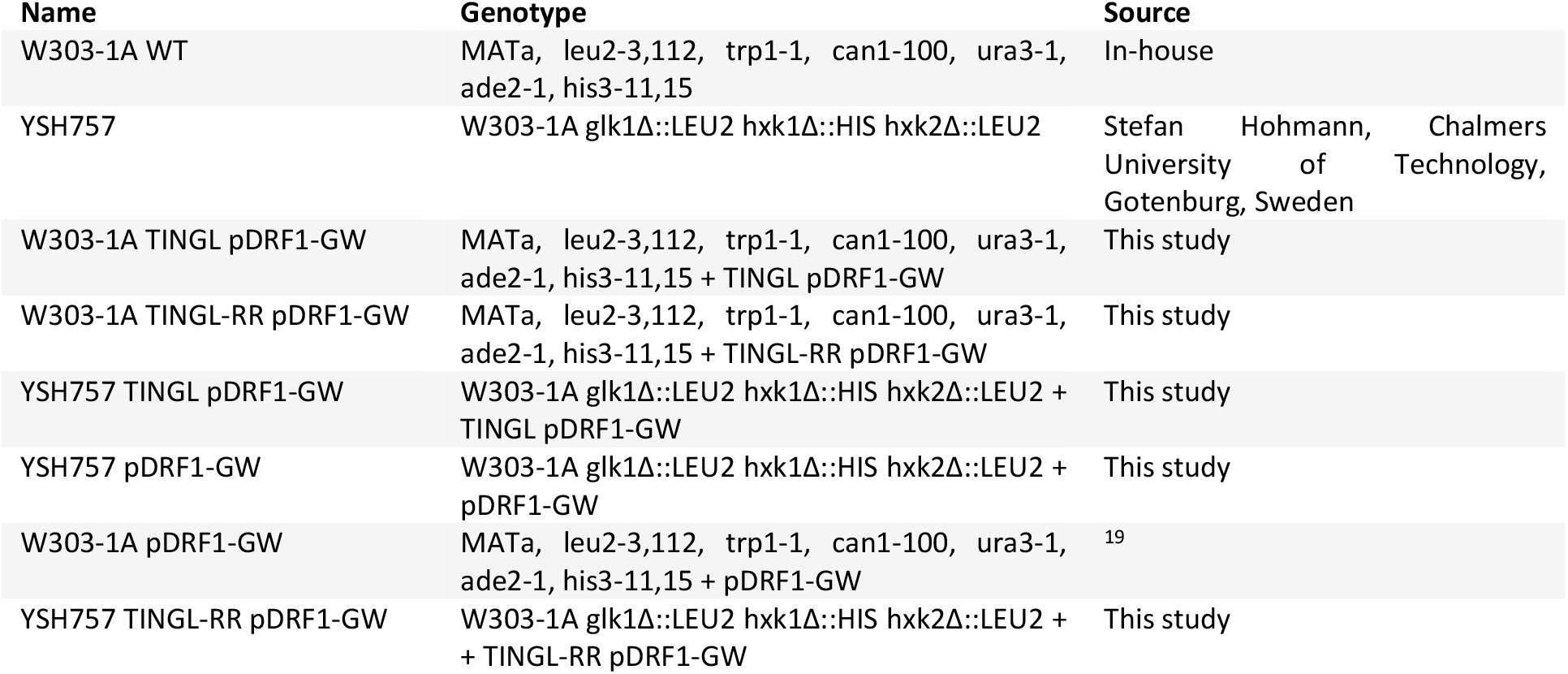
Yeast strains used in this study.

### Library generation

pYES2-ymTq2 (table 2). was amplified using ymTq2_146_FW and ymTq2_146_RV (listed in table 2). Mglb was amplified from Glifon600 (plasmid #126207) using primers muta_mglb146_1XNNK_FW and muta_mglb146_1XNNK _RV (listed in table 3) using KOD one polymerase (Toyoba, Osaka, Japan). Next, PCR products were digested using DpnI (NEB, Ipswich, MA) and purified using the MicroElute Cycle Pure kit (Omega, Norcross, GA). Thereafter, either a Gibson assembly (NEB) was performed according to manufacturer’s protocol followed by transformation, or samples were directly transformed in the YSH757 yeast cells according Gietz et al., 2007^20^. For the samples that were directly transformed, insert and vector DNA was added to the transformation mix in a 650 ng:100 ng vector:insert ratio. The transformed samples were plated on plates containing 1x YNB supplement with 2% agarose, 20 mg/L adenine hemisulfate, 20 mg/L L-tryptophan, 20 mg/L histidine-HCl, 60 mg/L L-leucine and 100 mM pyruvate.

**Table 2.**
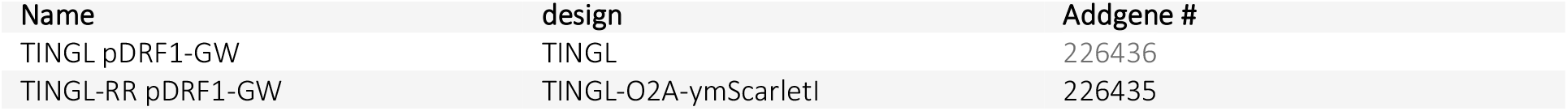
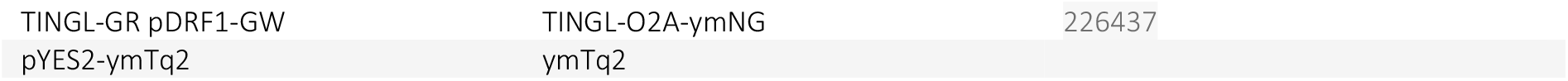
Used and made plasmids in this study.

| Name | design | Addgene # |
| --- | --- | --- |
| TINGL pDRF1-GW | TINGL | 226436 |
| TINGL-RR pDRF1-GW | TINGL-O2A-ymScarletI | 226435 |

**Table 3.**
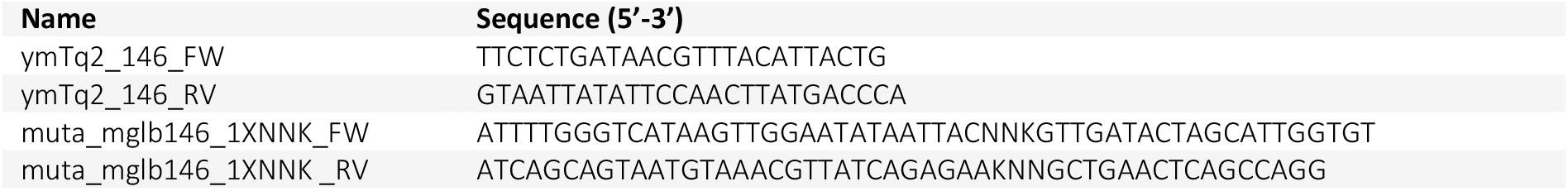
primers used to generate the sensor library.

### Library screening

Yeast cells expressing the glucose sensor libraries were visualized using a Amersham imager 600 (GE Healthcare, Chicago, IL, USA). First a fluorescence CFP image was taken after which a 1 M glucose solution was sprayed on the plates. Next, the plate was visualized again using the same settings as the first image. Differences between the two images were measured using an in-house Fiji macro^21^ which divides the two images and segments the colonies. The best performing colonies were selected and stored at -80°C and plasmid DNA was extracted using the Zymoprep Yeast Plasmid Miniprep Kits (Zymo Research, Irvine, CA, United States). Next, the sensor candidates were amplified using KOD one polymerase with 5’- ATCCCGGGAGGTCCTGGGTTGGATTCAACATCTCCACACAACTTCAATAGGTCGTAATTGGACTTACCAGAACCTTTATACAATTCATCCATACC-3’ as FW primer and 5’-TATGCTAGCATGGTTAGTAAAGGTGAAGAATTGTTC- 3’ as RV primer. Next, the products and a pDRF1-GW plasmid containing ymScarletI were digested using NheI and XmaI (NEB) and the products were ligated, yielding pDRF1-GW containing TINGL-O2A- ymScarletI (table 2). Next, ymNeongreen (ymNG) were amplified using 5’- TATCCCGGGATGGTCTCTAAGGGTGAAGA-3’ as FW and 5’-TTGTACAAGAAAGCTGGGT-3’ as RV primer. Next, the product and TINGL-O2A-ymScarletI were digested using XmaI and NotI (NEB) and the products were ligated in the plasmid, replacing ymScarletI with ymNG. Finally, TINGL was amplified using 5’-TATGCTAGCAAGCTTGAGCTCATCGATA-3’ and 5’-TCTAGATGCATGCTCGAG-3’ as FW and RV primers, respectively. Next, a pDRF1 plasmid and the PCR product were digested using NheI and NotI (NEB) and the TINGL sensor was ligated into pDRF1 which yielded TINGL pDRF1-GW.

### Dose response (plate reader)

YSH757 cells expressing either pDRF1-GW or pDRF1-GW TINGL-RR were grown on 2% agarose plates as described in library generation. Next, cells were resuspended in 1x YNB supplemented with, 20 mg/L adenine hemisulfate, 20 mg/L L-tryptophan, 20 mg/L histidine-HCl, 60 mg/L L-leucine and no carbon source to an OD of 2. Next, the 96-wells plate was filled with 90 μL of glucose at various concentrations after which 10 μL of cells was added. Fluorescence was measured using a Clariostar (BMG Labtech, Ortenberg, Germany) using 430/20 nm excitation and 480/20 nm emission for CFP signal and 570/15 nm excitation and 620/20 nm emission for RFP signal. Next, autofluorescence was subtracted using the signal from YSH757 expressing pDRF1-GW and the absolute TINGL signal as well as the ratio (TINGL signal divided by the RFP signal) was calculated. The values were normalized to the glucose concentration giving the lowest signal. Next, a dose-response curve was fitted according to equation 1 and equation 2 with Hillslope being the Hillslope, [glucose] the glucose concentration in mM, Kd the dissocation constant, top the maximal fluorescence value and bottom the minimal fluorescence value.

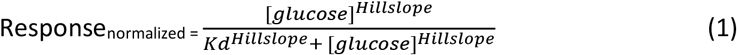

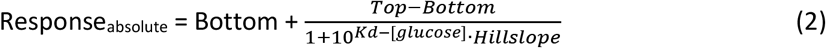

Finally, spectra were recorded as well using the Clariostar plate reader where emission spectra were recorded from 441 to 665 nm using a step size of 2 nm and a width of 10 nm with excitation set at 408/16 nm (429 nm LP dichroic). Next, fluorescence was normalized to the wavelength and glucose concentration giving the highest value using R version 4.4.1.

### Microscopy

W303-1A and YSH757 cells expressing either TINGL pDRF1-GW or TINGL-RR pDRF1-GW were grown 1x YNB containing 20 mg/L adenine hemisulfate, 20 mg/L L-tryptophan, 20 mg/L histidine-HCl, 60 mg/L L-leucine and either 100 mM glucose or 1% ethanol overnight at 30°C and 200 rpm after which the cells were diluted and grown again to mid-log (OD_600_ 0.1-1). Next, the cells were incubated in 6-wells plate containing ConA-coated coverslips (preparation described in^22^) and the coverslips were loaded in an attofluor cell chamber (Thermo Fisher Scientific, Waltham, MA, USA) after which 1 mL of medium was added. Coverslips were loaded in the stageholder of a Nikon Ti-eclipse widefield fluorescence microscope (Nikon, Minato, Tokio, Japan) set at 30 °C. Fluorescence was obtained using 438/24 nm excitation, 458LP dichroic and 483/32 nm emission for CFP signal and using 570/20 nm, 600LP dichroic and 610LP emission for RFP signal. Fluorescence was recorded using an Andor Zyla 5.5 sCMOS Cameras (Andor, Belfast, Northern Ireland) and a SOLA 6-LCR-SB power source (Lumencor, Beaverton, OR, USA). For the various timelapse movies, a 3 or 5 minute baseline was recorder after which the compound of choice was added to the desired concentration. For the dose-response, subsequent pulses of glucose was added to YSH757 expressing TINGL-RR to set the glucose concentration at various concentrations. Washing and starvation was performed in the attofluor cell chamber, where cells were rinsed twice with 1 mL of growth medium where the carbon source was omitted. Next, cells were incubated for 15- 20 minutes and visualized as described. Data analysis was performed using R version 4.4.1.

Fluorescence lifetimes were measured by growing YSH757 cells expressing TINGL on plates containing 1x YNB supplement with 2% agarose, 20 mg/L adenine hemisulfate, 20 mg/L L-tryptophan, 20 mg/L histidine-HCl, 60 mg/L L-leucine and 100 mM galactose. Next, the cells were resuspended 1x YNB containing 20 mg/L adenine hemisulfate, 20 mg/L L-tryptophan, 20 mg/L histidine-HCl, 60 mg/L L-leucine and either 0 or 10 mM glucose. Cells were transferred to a 24 wells plate, incubated for 15 minutes on room temperature, and visualized using a Leica Stellaris8 laser scanning confocal microscope (Leica, Wetzlar, Germany), equipped with White Light Laser and using a HC PL APO 40x/0.95 DRY lens. Pinhole size was set to 1AU. Samples were illuminated with 448 nm excitation light set at a frequency of 40 MHz. Images with 512 x 512 pixels, pixel size 569 nm, were acquired using a HyDX2 detector in photon counting mode. The mTurquoise2 acquisition parameters were: emission wavelength range of 460-520nm, dwell time 2.82 μs, line averaging of 8, imaging frequency of 400 Hz. The accompanied software was used to determine intensity weighted lifetimes of a 2-component decay fit.

### Growth assay

W303-1A cells expressing pDRF1-GW, TINGL pDRF1-GW and TINGL-RR pDRF1-GW were grown as described in the microscopy section. Cells were washed and concentrated to an OD_600_ of 1 with the same medium with carbon source omitted. Afterwards, cells were transferred a final OD_600_ of 0.04 in a 48-multiwell culture plates containing 480 μL of fresh medium with either 0.1% ethanol, 10 mM galactose or 10 mM glucose. Cells were grown in the Clariostar plate reader at 30°C and 700 rpm orbital shaking. OD_600_ was measured every 5 minutes for 25 hours with 30 flashes per cycle. Data analysis was performed using R version 4.4.1. In brief, OD_600_ values were corrected for empty medium absorbance and a smoothing average was performed. A sliding window was performed where a linear fit was performed on the ln-transformed data. This slope indicates the growth rate for every window. Lag phase was determined as the time needed to obtain the maximal growth rate.

### Cell free extracts and dose-response curves at various pH

YSH757 cells expressing pDRF1-GW, TINGL pDRF1-GW pDRF1-GW were grown overnight at 200 rpm and 30°C in 2x YPAgal medium containing 20 g/L bacto yeast extract, 40 g/L bacto peptone, 160 mg/L adenine hemisulfate and 40 g/L galactose or in 2x YNB containing all 152 mg/L of all standard amino acids (with leucine at 380 mg/L), 2% galactose and 2% pyruvate. Subsequently, cell cultures were centrifuged at 14.000 g for 5 min at 4°C. Supernatants were discarded and cells were washed three times in 4°C freeze buffer containing 0.01M KH_2_PO_4_ with 0.75 g/L EDTA. Next, the cell suspensions were centrifuged at 14.000 g for 1 min at 4°C and supernatants were discarded. Cell pellets were resuspended in ice-cold sonification buffer containing 0.1M KH_2_PO_4_ with 0.41 g/L MgCl_2_ and centrifuged again at 14.000 g for 1 min at 4°C. Next, supernatants were discarded, and cell pellets resuspended in ice-cold sonification buffer with 10 mM DTT. Cell suspensions were transferred to a pre-cooled screw cap tube with 0.75 grams of glass beads. Cells were shaken in 8 bursts of 10 s at speed 6 in a FastPrep® homogenizer (MP bio, Santa Ana, CA) with cooling the suspension for at least 60 s on ice between bursts. Afterwards, the cell suspensions were centrifuged at 14.000 g for 15 minutes at 4°C. Supernatants were transferred to a fresh pre-cooled tube, snap frozen and stored at - 80°C. Protein concentrations were determined using the Pierce™ BCA Protein Assay Kit (23225, Thermo Scientific, Waltham, MA) following the manufacturer’s instructions for the microplate procedure. OD_562_ of the standards and samples in duplicates were measured using the multiskan go plate reader (Thermo Fisher Scientific). Data analysis was performed using Excel. OD_562_ was corrected for auto-absorbance of the blanks. Protein concentrations were interpolated using the TREND function. Next, the extracts were diluted 10x in PBS at pH 6.3, 6.5, 7.0. and 7.5 containing various glucose concentrations. CFP and RFP signal was obtained using 430/20 nm and 570/15 nm excitation, 455 and 594 nm LP, 480/20 nm and 620/20 nm emission for CFP and RFP, respectively. Next, the fluorescence signal was corrected for background signal using fluorescence of extracts of YSH757 cells expressing the empty pDRF1 plasmid and CFP:RFP ratios were calculated. Lastly, a Hillfit was performed according equations 1 and 2 in R version 4.4.1.

### Response speed assay

90 μL of cell-free extracts were loaded inot a 96-well plate. Next, fluorescence was measured before and after pulsing glucose using a FLUOstar Omega microplate reader (BMG Labtech) with a 430/10 nm excitation filter and 480/10 nm emission filter, top optics, and a gain of 1500. Two kinetic windows were used. Kinetic window 1 with 10 cycles of 1 s, measurement starting time 0.0 s and 15 flashes per well per cycle. Consequently, glucose to an end concentration of 100 mM or demineralized water was added with a pump speed of 300 μL/s. Afterwards, kinetic window 2 started with 120 cycles of 1 s, measurement starting time 10.5 s, and 15 flashes per well per cycle. Data analysis was performed using GraphPad prism 9 software. A curve was fitted through the datapoints using equation 3 with the plateau being asymptote, K the rate constant, Y_0_ the starting fluorescence and X the time.

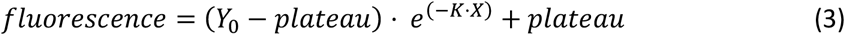

### THINGL development

THINGL and THINGL-RR were generated by performing a PCR using KOD one polymerase on the mglb domain of TINGL using 5’-ATGGTTGATACTAGCATTGGTGTAAC-3’ and 5’-GTAGCTGAACTCAGCCAGG-3’ as FW and RV primers, respectively. Simultaneously, a PCR with KOD one polymerase was performed on mScarletI-T2A-mTq2 in pDRF1 using 5’- GACAACCTGGCTGAGTTCAGCTACTTTAGCGACAACGTCTATATCAC-3’ and 5’- GTTACACCAATGCTAGTATCAACCATGTAGTTGTACTCCAGCTTG-3’ as FW and RV primers, respectively. Next, a Gibson assembly was performed which yielded mScarletI-T2A-THINGL (THINGL-RR) in pDRF1. Next, a PCR was performed on THINGL-RR using either primers 5’- TATAAGCTTATGGTGAGCAAGGGC-3’ and 5’-GCTTCTAGATTACTTGTACAGCTCGTCC-3’ or primers 5’-TATAAGCTTATGCGGTTCAAGGTGC-3’ and 5’-GCTTCTAGATTACTTGTACAGCTCGTCC-3’. Next, the PCR products and a pcDNA3.1 vector were digested using HindIII and XbaI (New England Biolabs, Ipswich, MA) after which THINGL-RR and THINGL were ligated into pcDNA3.1, which yielded THINGL-RR (mScarletI-T2A-THINGL) and THINGL in pcDNA3.1.

### Cell Culture and Transfection

HeLa cells (CCL-2, American Type Culture Collection) were cultured in Dulbecco’s modified Eagle medium supplemented with GlutaMAX (DMEM + GlutaMAX, 61965, Gibco), 10% fetal bovine serum (FBS, 10270, Gibco), and maintained at 37 °C in a humidified atmosphere containing 7% CO_2_. No antibiotics were added to the culture medium at any stage. For transfection, a DNA–PEI complex was prepared in Opti-MEM. After incubation for 20 minutes at room temperature, the transfection mixture was added directly to the cells.

### Sample Preparation for Imaging

Twenty-four hours after transfection, the culture medium was replaced with microscopy medium lacking glucose, composed of 137 mM NaCl, 5.4 mM KCl, 1.8 mM CaCl_2_, 0.8 mM MgSO_4_, and 20 mM HEPES, adjusted to pH 7.4. Cells were incubated in this glucose-free medium for at least 10 minutes at room temperature prior to imaging. During live-cell imaging, 5 μL of a 1 M glucose stock solution was added directly to the imaging chamber, resulting in a final glucose concentration of 5 mM.

### Fluorescence Lifetime Imaging (FLIM)

FLIM measurements were performed with excitation at 448 nm and a modulation frequency of 40 MHz. Emission was detected in the 460–550 nm range using a HyD-X2 detector operated in photon counting mode. Images were acquired at 512 × 512 pixels, covering a field of view of 290 μm × 290 μm. To improve the signal-to-noise ratio, eight line repetitions were used. The scanning speed was set at 400 Hz, corresponding to a pixel dwell time of 2.8 μs and an acquisition time of 10.32 seconds per frame. Time-lapse series were recorded at intervals of 15 seconds between frames. The pinhole diameter was set to 3.25 Airy units, and a zoom factor of 1 was applied.

FLIM datasets were exported as TIFF image stacks compatible with ImageJ/Fiji software, with photon counts scaled such that each grey level represented one count within the 16-bit dynamic range.

Fluorescence lifetime values were mapped linearly from 0 to 10 nanoseconds across the full 16-bit greyscale range. In Fiji, regions of interest (ROIs) were manually defined around individual cells, the data was exported as CSV files and further process in R. Full details are reported here: https://github.com/JoachimGoedhart/Simple-lifetime-timelapse-analysis/.

## Results

### Library screening yields a responsive glucose sensor

To generate a pH independent biosensor we started with mTq2 as the fluorescent protein of choice since this FP is rather pH insensitive ^23,24^. Using the same design principles as for GFP-based single-FP biosensors^25^ we inserted the glucose binding domain MglB inside ymTq2 at position 146 with 1 random amino acid at each side as a linker. The libraries were generated by performing a PCR on ymTq2 in the pYES2 expression plasmid (linearized at amino acid 146 of ymTq2) and the binding domain with the random amino acid on each side plus 20 basepairs homology to ymTq2. Next, a Gibson assembly was performed, or the two PCR products were directly transformed into yeast. We choose to use YSH757 as the yeast strain of choice since this strain lacks the enzymes *HXK1, HXK2 and GLK1* that convert glucose into glucose-6-phosphate (and therefore cannot metabolize it) meaning that intracellular glucose equilibrates with the extracellular glucose concentration. Next, the cells were plated on plates containing 100 mM pyruvate as the carbon source and CFP signals of the plate were recorded using a fluorescence western blot imager. Thereafter, a 1 M glucose solution was sprayed on the plate and the plate was recorded again using the same settings (figs. 1A and 1B). In this way we could conveniently screen the responsiveness of sensor candidates *in vivo*. Colonies that changed the most in CFP signal were picked up and further characterized. From these candidates only one variant (variant 5) turned out to be robust in terms of monomericity, brightness, specificity and response speed. This variant contained a methionine and tyrosine as the 2 linkers (fig. 1C). We named this sensor TINGL (Turquoise INdicator for GLucose). Since the sensor was not ratiometric by itself (fig. 1D), we combined it with ymNeongreen or ymScarletI using the polycistronic element O2A to generate loose expression of the sensor with these FPs. These constructs were named TINGL-GR (TINGL-Green Ratiometric) and TINGL- RR (TINGL-Red Ratiometric). These 2 constructs can be used to normalize the signal of TINGL. Next, using the glucose phosphorylation deficient YSH757 strain we were able to make *in-vivo* dose-response curves of TINGL and TINGL-RR in the plate reader by adding glucose to various concentrations in a 96- wells plate harboring these two strains (fig. 1E). The dose-response curves gave a K_d_ of 1.8 and 1.1 mM and a Hill slope of 0.8 and 0.6 for TINGL and TINGL-RR, respectively. The sensor is also rather pH-robust down to a pH of 6.5. Below this value the CFP:RFP ratios are largely affected by pH (fig. 1F). Furthermore, we were able to assess the response rate of the sensor by pulsing glucose to a final concentration of 100 mM to cell-free extracts of YSH757 expressing TINGL (fig. 1G). The response kinetics (one-phase decay) showed a half-time of 4.9 seconds. Fluorescence lifetime of the sensor in YSH757 cells changed with 0.3 ns when stimulated with 10 mM glucose (fig. 1H). The sensor showed a uniform distribution within the cells (fig. 1H), which was not the case for other sensor variants. Lastly, we performed a growth assay of W303-1A cells expressing TINGL-RR to test for any burden on the cells. We found for some conditions a very small, yet significant (Wilcoxon test, α = 0.05), effect of the sensor. Only on galactose the cells had a slight reduced growth rate (0.23 h^-1^ for TINGL-RR and 0.24 h^- 1^ for the empty plasmid) and the lag phase was slightly increased for both glucose (by 0.7 hours) and galactose (by 1.6 hours) as substrate. The maximal OD_600_ (used as a proxy for yield) was minimally changed for glucose as substrate only (0.45 for cells expressing TINGL-RR and 0.46 for cells expressing the empty plasmid). In conclusion, TINGL is a robust, functional sensor suitable for measuring glucose levels inside single cells.

**Figure 1.**
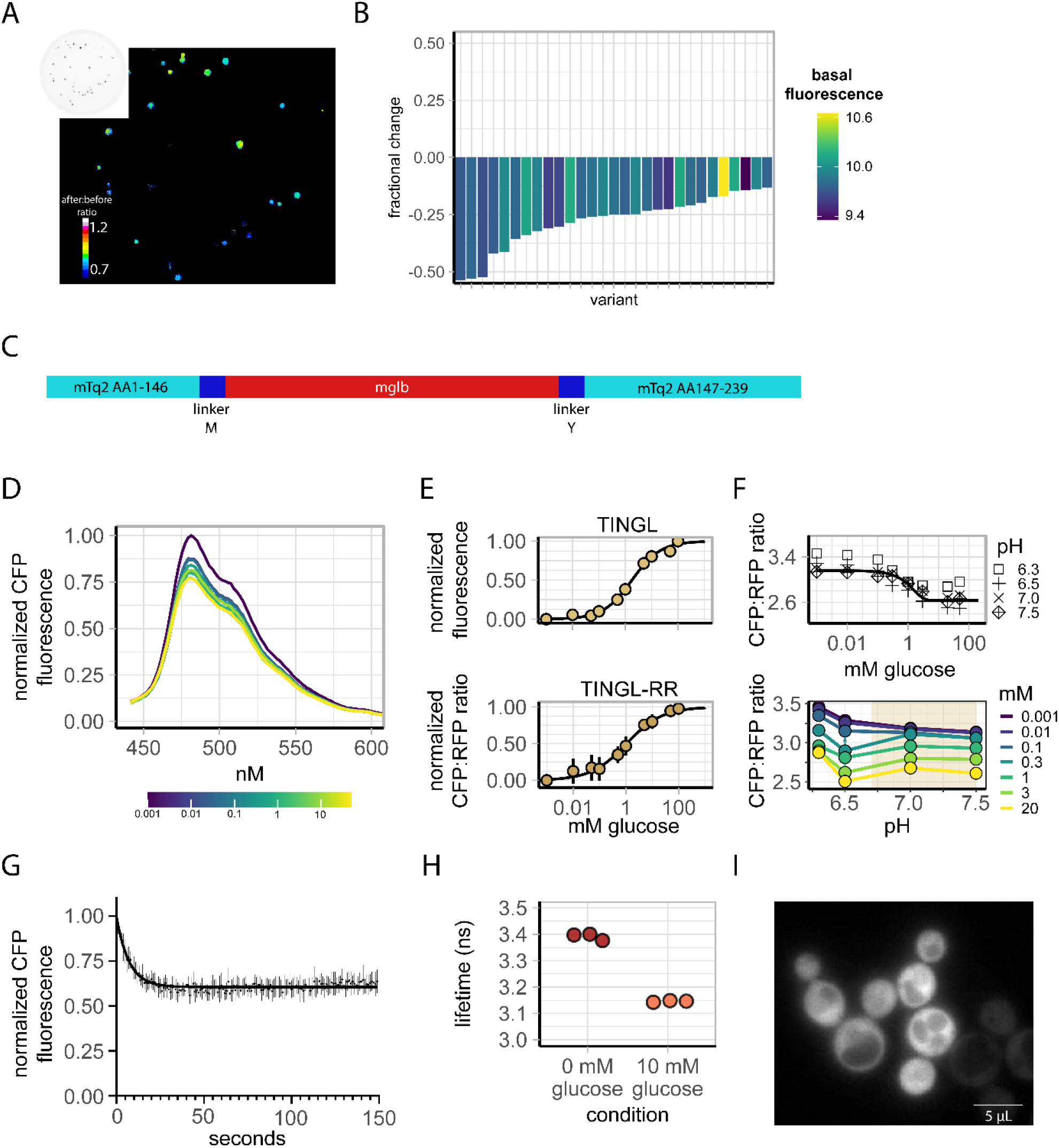
Screening and characterization of glucose sensors. A) The screening assay identifies responsive glucose sensors in a medium-throughput manner. Plates lacking glucose have YSH757 yeast cells expressing sensor variants and are visualized whereafter glucose is sprayed on the plate and visualized again. Dividing the 2 images yields a ratiometric image showing which colonies have a responsive sensor. Colour indicates ratio of fluorescence after the spray divided by the fluorescence before the spray. B) normalized change of each colony tested on the plate, colour indicates basal fluorescence. C) design of the best performing variant. D) In-vivo CFP emission spectrum of YSH757 cells expressing TINGL. Colour indicates amount of glucose added to the cells. Data obtained in a fluorescence plate reader E) In-vivo Dose-response curve of YSH757 cells expressing TINGL or TINGL-RR obtained using a plate-reader. Fluorescence and CFP:RFP ratio are in absolute values. Line shows a Hillfit through the data points. F) Dose-response curve of TINGL-RR of cell-free extracts. Top facet shows CFP:RFP ratio versus mM glucose with the shapes depicting the pH. Line shows a Hillfit through the data of pH 7.5, 7 and 6.5 combined. Colored panel indicates the physiological pH of yeast. G) In-vitro response speed of the sensor. Cell-free extracts YSH757 cells expressing TINGL were pulsed with glucose to a final concentration of 100 mM glucose at t=0 seconds. H) lifetimes of YSH757 yeast cells expressing TINGL in medium containing YNB medium with either 0 mM or 10 mM glucose, depicted by the colour of the points. Each points represents a biological replicate. I) example of the sensor distribution in W303-1A cells.

### TINGL is a glucose-specific biosensor

Although the initial characterization (fig. 1) were promising, it is unclear whether the biosensor is fully functional in cells during dynamic conditions. Furthermore, the specificity of the sensor should also be tested. We performed a variety of transitions for W303-1A cells expressing TINGL or TINGL-RR to test the robustness and specificity of the sensor. We grew cells on 1% EtOH, visualized them using a widefield fluorescence microscope and added 10 mM fructose after three minutes and 10 mM glucose after seven minutes (fig. 2). Fructose gives an almost identical, even a slightly stronger startup of glycolysis compared to glucose based on pH readouts (data not shown), implying that all intracellular changes after a glucose pulse also occur when fructose is added, except for the presence of glucose itself. We found no change in CFP signal after the fructose addition (normalized CFP/RFP ratio of 0.99 ± 0.04, fig. 2A) whereas glucose addition gave a decrease of more than 20%, showing that TINGL is specific for glucose. We found the same result for mannose, (normalized CFP/RFP ratio of 0.97 ± 0.07, fig. 2B). Next, we tested how sucrose (a dimer of glucose and fructose) addition affects intracellular glucose levels. We found a very small response after a 0.3 mM sucrose pulse whereas an additional pulse of 10 mM sucrose gave a response to normalized ratio of 0.88 ± 0.05, which is less of a strong response than a glucose pulse of 10 mM. This was expected as sucrose gives combinatorial (competitive) import of both glucose and fructose by the same hexose transporters. Next, we wondered if a 0.3 mM glucose pulse would give a measurable change in intracellular glucose levels since the K_50_ of the glucose-sensing system cAMP-PKA is approximately 0.3 mM and this is also the concentration where yeast cells start to ferment^26^. We indeed found a small and transient response of the sensor after the 0.3 mM glucose addition, also showing that the sensor picks up these small changes in glucose concentrations. The second addition of 10 mM glucose resulted again in a response of 20% of the sensor. Lastly, we measured glucose levels for a longer period of time after a 10 mM glucose pulse. Interestingly, after 20 minutes glucose levels started to increase over time, indicating changes in the activity of glycolysis and/or glucose transport, through post-translational modifications or gene expression. In summary, TINGL can be used conveniently to robustly measure glucose dynamics during various transitions.

**Figure 2.**
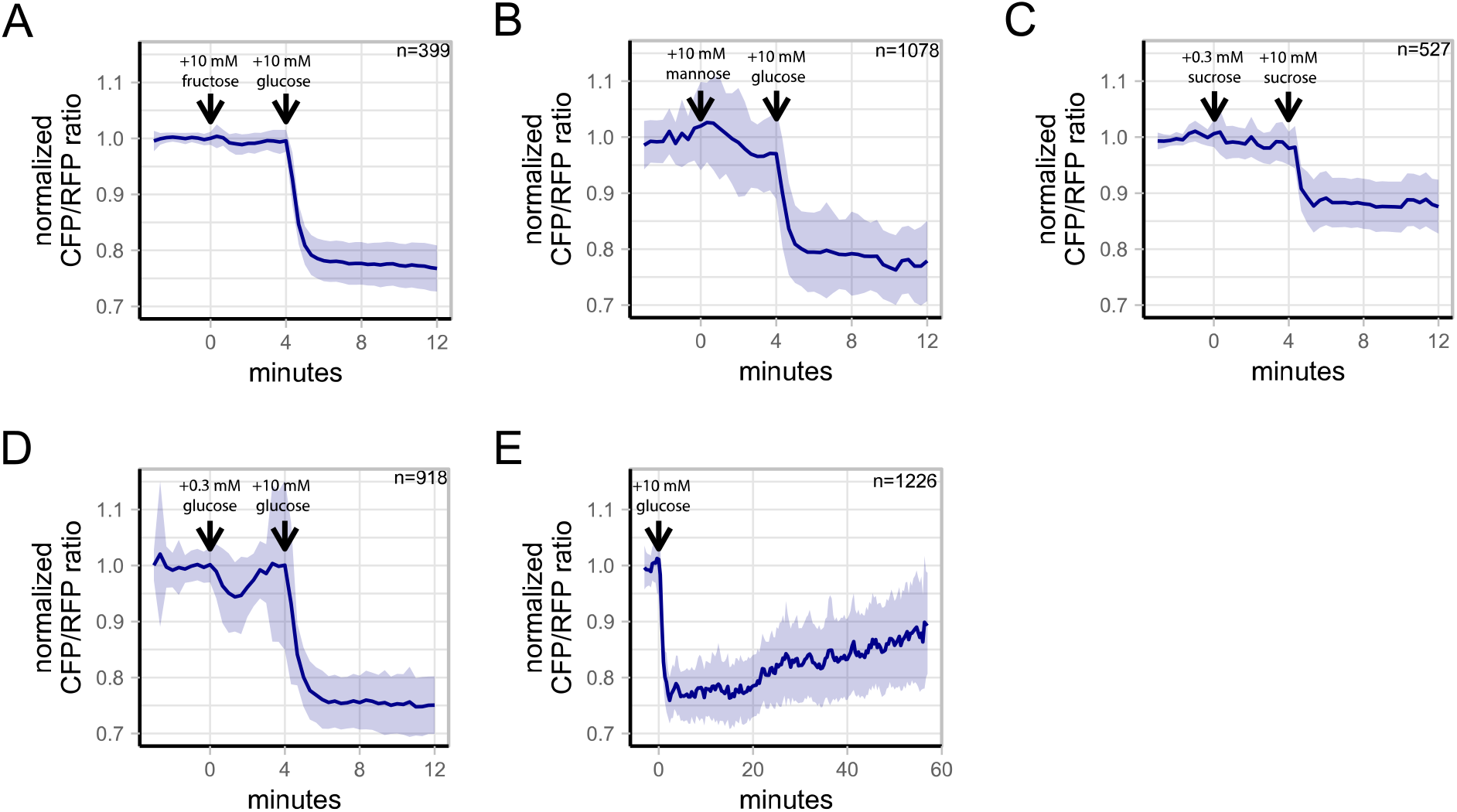
TINGL-RR responses during various transitions show that the sensor is a specific and robust sensor. A) Normalized CFP/RFP ratios of TINGL-RR of W303-1A cells grown on 1% EtOH to a 10 mM fructose and glucose pulse. B). Normalized CFP/RFP ratios of TINGL-RR of W303-1A cells grown on 1% EtOH to a 10 mM mannose and glucose pulse. C) Normalized CFP/RFP ratios of TINGL-RR of W303-1A cells grown on 1% EtOH to a 0.3 and 10 mM sucrose pulse. D) Normalized CFP/RFP ratios of TINGL-RR of W303-1A cells grown on 1% EtOH to a 0.3 and 10 mM glucose pulse. E) Normalized CFP/RFP ratios of TINGL-RR of W303-1A cells grown on 1% EtOH to a 10 mM glucose pulse. Lines indicate mean CFP:RFP ratio, normalized to the baseline. Shades indicate SD. Arrows indicate addition of the sugar, stated above the arrow. Data from three biological replicates.

### A hexokinase NULL mutant can be used to decipher the absolute internal glucose concentration

Fluorescent biosensors are difficult (almost impossible) to use for absolute concentration measurements in living cells. This is because dose-response curves need to be made which is done *in vitro* for most sensors and these conditions do not resemble the actual cellular conditions (and visualization setup) in which the sensor is eventually used^23,27^. However, because glucose transport in yeast is mediated by carrier-mediated diffusion, we can make dose-response curves *in vivo* using the YSH757 yeast strain. In this strain, the intracellular glucose level will equilibrate with the extracellular glucose concentration. In principle, this dose-response curve can be used to decipher intracellular glucose concentrations. Using the dose-response of the hexokinase NULL-mutant, we were able to translate the CFP:RFP ratio to actual intracellular glucose concentrations (figs. 3 and S2). We found that a 10 mM glucose pulse to de-repressed yeast cells results in an intracellular glucose concentration of approximately 1 mM, which is in line with previous results based on biochemical assays^5^. As already shown in fig. 2E, glucose concentrations drops again to approximately 0.5 mM after 20 minutes. Thus, using TINGL, intracellular glucose concentrations can be determined in single cells in time.

**Figure 3.**
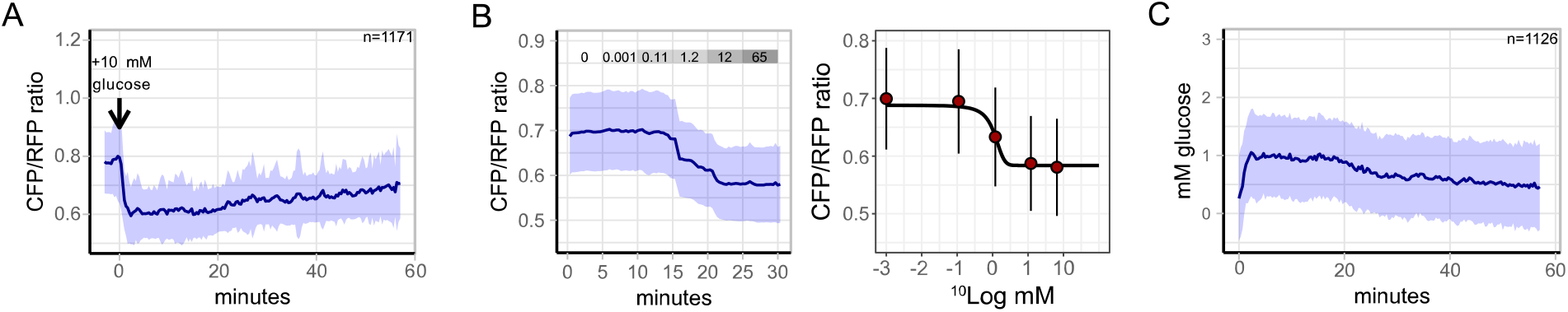
TINGL-RR can be used to measure absolute glucose concentrations using the hexokinase mutant. A) CFP/RFP ratios of W303-1A cells grown on 1% EtOH to a 100 mM glucose pulse. Lines show the mean CFP/RFP ratio, shades indicate SD. B) In-vivo dose-response curves using the hexokinase mutant YSH757. Grey bars depict the concentration of glucose, in mM, in the medium. Blue lines show mean CFP/RFP ratio, shades indicate SD. Black line shows the Hillfit curve, dots and whiskers indicate mean CFP/RFP ratio with SD, respectively. C) The obtained dose-response curve can be used to calculate the absolute glucose concentrations in cells. Note that the number of cells is different compared to panel A. This is because cells can have ratios outside the dose-response curve. This results in infinite values which cannot be depicted. The n values shown here are therefore the total number of cells, but at single timepoints the number of cells that show measurable CFP/RFP ratios can be lower. Lines show mean CFP/RFP ratio, shades indicate SD. Data obtained from three biological replicates.

### Glucose repression affects the intracellular glucose dynamics during a glucose transition

Lastly, we tested how glucose repression affects the response to glucose (re)addition. Glucose repression is known to affect the maximal glycolytic flux and to change the composition of the hexose transporters^7,28–30^. We tested the initial glucose transport dynamics for cells that were glucose repressed or de-repressed (fig. 4). Correspondingly, W303-1A cells were grown on either 100 mM glucose or 1% EtOH and washed twice with media containing no carbon source. Next, the cells were visualized on the microscope and 10 mM or 100 mM glucose was pulsed after 3 minutes. Interestingly, cells pulsed with 10 mM glucose showed a somewhat more dynamic response compared to cells pulsed with 100 mM glucose (fig. 4), although this effect was small. Trajectories stabilized after approximately 10 minutes where we found that cells pre-grown on glucose showed a higher normalized CFP:RFP ratio compared to cells grown on EtOH, meaning that the glucose level is lower (Student’s t-test, p < 0.01, fig. 4A, 4B). This is in line with the data in fig. 3, where we showed that cells that adapt to a transition from ethanol to 10 mM glucose start to decrease intracellular glucose concentration after 20 minutes, where we suspect glucose repression starts to kick in. In conclusion, glucose repression does affect the cellular response to a glucose addition as they obtain lower intracellular glucose concentration compared to de-repressed cells.

**Figure 4.**
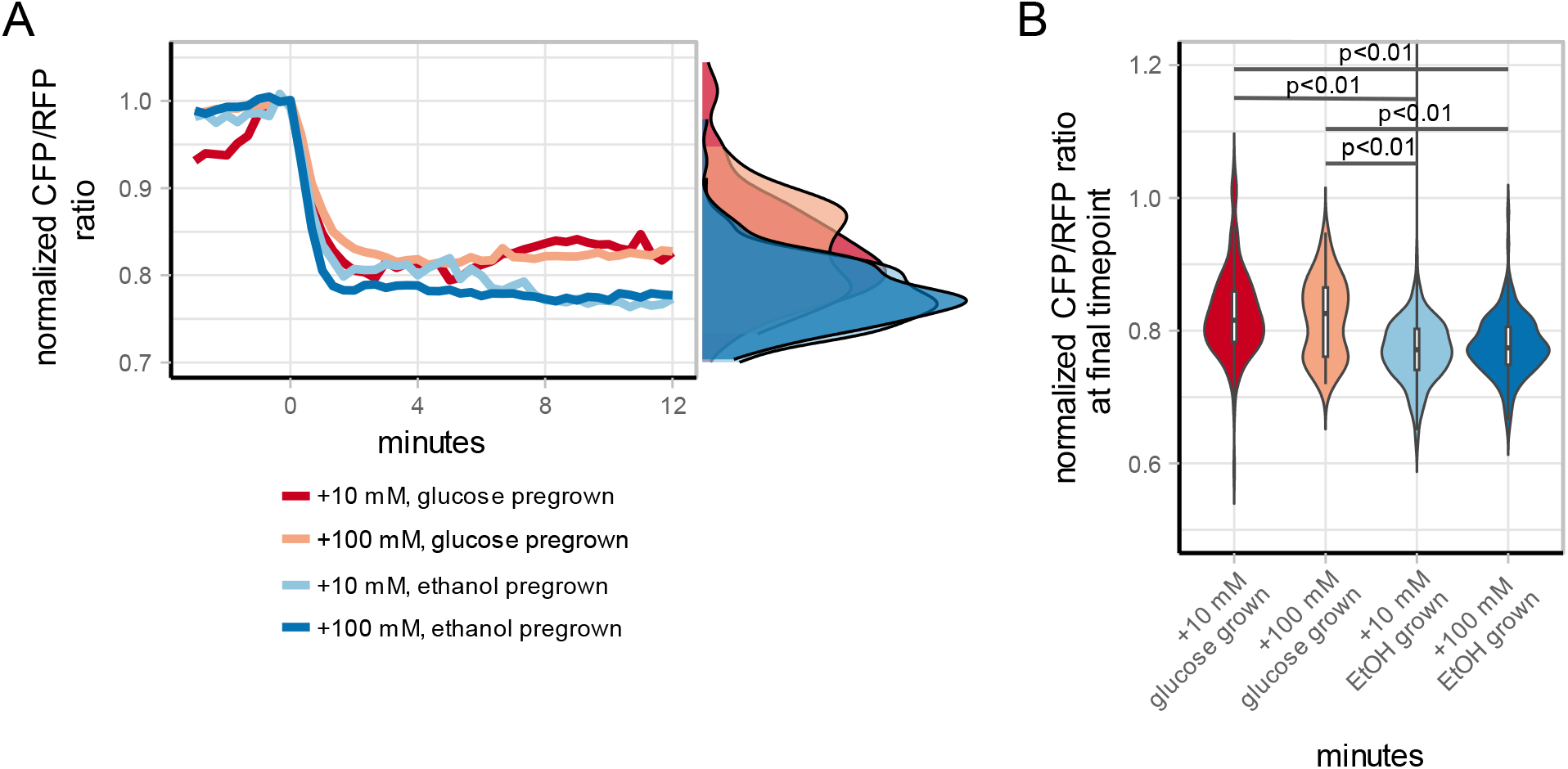
A glucose transition gives different dynamics, based upon glucose repression state. A) normalized mean CFP:RFP ratios of TINGL-RR of W303-1A cells during a transition to either 100 or 10 mM glucose. Cells were pregrown on either 1% EtOH or 100 mM glucose, washed in medium without carbon source, loaded into the sample holder whereafter glucose was pulsed at 0 minutes. The right side shows a normalized distribution of the 4 transition at t = 12 minutes. Colours indicate transition. B) Violin plot, together with a boxplot depicting the maximal decrease of CFP:RFP ratio (i.e. the maximal amount of glucose in a cell) of the 4 transitions. Boxplots indicate median with quartiles; whiskers indicate largest and smallest observations at 1.5 times the interquartile range. p values of Student’s t-test are shown for the significant differences. Data obtained from at least 3 biological replicates.

### The human codon-optimized version, THINGL, robustly indicated glucose levels in HeLa cells

In addition to yeast cells, a robust cyan glucose sensor would also be useful for research in human cells. Therefore, we decided to copy the TINGL design into human codon-optimized mTurquoise2 and move the biosensor to an expression vector for human cells. HeLa cells were transfected with THINGL and starved for glucose for at least 10 minutes after which 5 mM glucose was added. CFP fluorescence as well as lifetimes of THINGL were measured for 4 replicates (fig. 5). Addition of glucose showed a similar response a was seen in yeast cells with a decrease in fluorescence up to 30%. Lifetimes decreased from 3.5 to 3.0 ns, which is a slightly larger decrease than found in yeast cells (fig. 1H). In conclusion, THINGL can be conveniently used to robustly measure changing glucose levels in living human cells.

**Figure 5.**
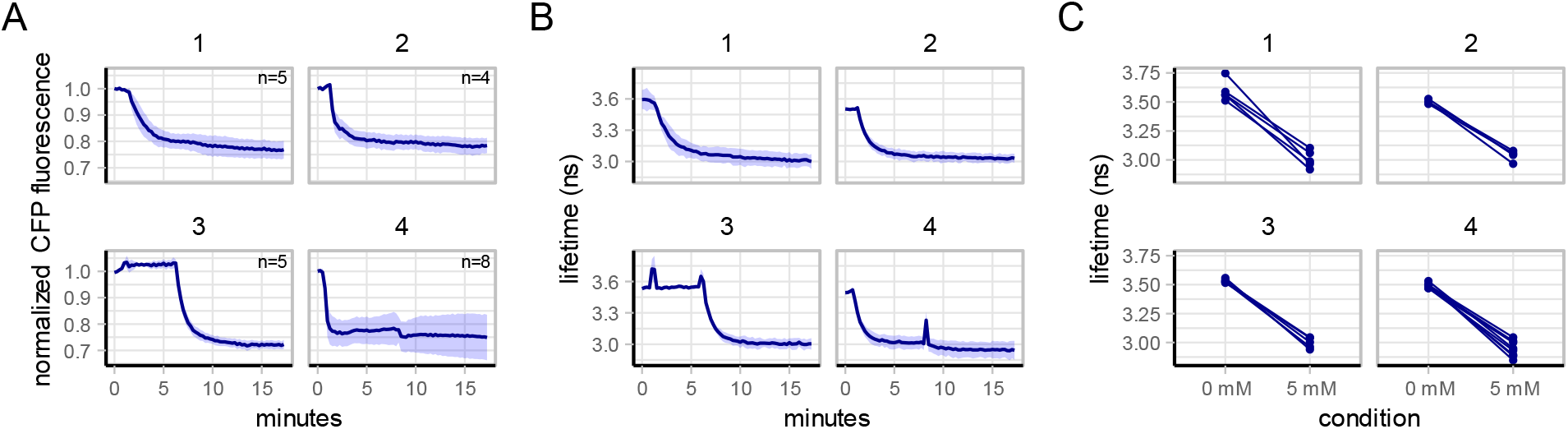
THINGL visualizes glucose levels in HeLa cells. A) Normalized CFP fluorescence signal of HeLa cells expressing THINGL during the transition to glucose in glucose-starved HeLa cells. Lines show mean fluorescence value, normalized to the first 3 timepoints, shades indicate SD. B) Fluorescence lifetimes of THINGL during the transitions. Lines show mean fluorescence value, normalized to the first 3 timepoints, shades indicate SD. C) The lifetimes of each cell for both conditions. Each point depicts a cell. Each facet depicts a replicate.

## Discussion

To study the dynamics of glucose in living cells, several glucose biosensors have been reported^8–11,13,18^. However, the currently available biosensors are pH sensitive, which is a major drawback for use in yeast, where the pH may drop to 6.5 during nutrient transitions. A Turquoise-based glucose biosensor that was reported while our work was in progress, showed a largely pH independent response, but the affinity of that biosensor was roughly one to two orders of magnitude too high for practical applications [ref]. Therefore, to enable glucose sensing in cells, we have developed a robust biosensor with an affinity that is compatible with intracellular glucose concentrations.

To achieve this, we have developed an efficient method to generate and screen for metabolic sensors (and perhaps also other sensors), in this case for glucose. Budding yeast is genetically versatile, which we exploited to generate sensor libraries; the transformed colonies on the plates can be sprayed by an inducer for the ligand of the sensor and screened for responsive candidates. In principle, this method can be used to screen biosensors for any ligand that can be induced in a yeast strain. In terms of glucose metabolism, this screening method can for example be used to develop biosensors for fructose-1,6-bisphosphate (an important flux-associated glycolytic intermediate^31–34^) or for glucose- 6P, which is an important branch point between glycolysis, glycogen (and trehalose) metabolism and the pentose phosphate pathway.

Using this method, we developed an mTurquoise2-based glucose sensor, named TINGL, which proves to be a robust and specific sensor for this metabolite. Using the assays in fig. 2 we found that TINGL is specific, robust and can measure low glucose concentrations. Furthermore, the sensor showed to be fast responding with a half-time of less than 5 seconds and is monomeric. The sensor did give small effects on cellular growth, although this was minimal; a reduced growth rate for galactose, a decrease in maximal OD_600_ for glucose as substrate and an increased lag phase for both. Although these effects were very small, this still raises a concern for sensors characterized in other systems than yeast where effects on cellular health are more difficult to measure. Sensors developed and tested in mammalian cells, for example, are often not tested for toxicity as growth rate determination is very complex for this organism. Therefore, we believe that the testing of these sensors in yeast can be beneficial for sensor development in general. The expression of TINGL and TINGL-RR using pDRF1 as a plasmid (which is a high expression plasmid) generates a surplus of signal from the sensor. Therefore, the growth defects can most likely be eliminated by using a lower expression plasmid if needed.

We exploited the genetic versatility of budding yeast further by performing an *in vivo* dose-response of TINGL-RR and used these data to translate the CFP:RFP ratios to actual glucose concentrations. Although this gives a loss of data (as cells can have fluorescence values outside the calibration curve) we still obtained enough data to measure actual dynamic glucose concentrations (fig. 3). An addition of 10 mM glucose to derepressed cells resulted in an average intracellular glucose concentration of 1 mM. Interestingly, we found a clear and consistent change in glucose levels after 20 minutes. This is in line with previous data, which also shows a clear change for some glycolytic intermediates after 20 minutes^35^. This is probably caused by a gradual transition to a glucose-repressed state of the cells, which results in a reprogrammed glycolytic state with an increase glycolytic flux^35,36^. The reprogramming of glycolytic enzymes, which includes transporters that have a lower affinity but higher capacity^7,29,37^, is known to increase glycolytic flux. This results, based on our data, in an apparent and perhaps somewhat counterintuitive decrease of intracellular glucose. With our glucose sensor at hand, it will be interesting to further investigate the potential role of intracellular glucose in glucose signalling and repression.

We also measured glucose dynamics of 15-minutes starved cells pre-grown on either ethanol or glucose to a glucose addition of either 10 or 100 mM. Although glucose-repressed cells express transporters with a K_m_ up to 100 mM (compared to 1-2 mM for de-repressed cells)^7^, we did not find a clear difference in the initial glucose dynamics. Yet, after the steady state was reached, we did find that glucose-repressed cells have lower glucose levels compared to repressed cells (fig. 4A). This must be caused by the fact that cells pre-grown on glucose were adapted to glucose growth and could obtain a higher glycolytic flux compared to cells grown on ethanol which is also in line with the glucose adaptation observed after 20 min in ethanol-grown cells (fig. 3).

Lastly, we show that THINGL, the glucose sensor for use in human cells shows a reproducible and robust response in both fluorescence intensity and lifetime after a glucose addition to starved HeLa cells. Thus, the design uncovered through the yeast screen is readily transferable to human cells, enabling the study of glucose dynamics and this screening method can be potentially used for the development of other sensors as well.

In conclusion, TINGL is a robust biosensor to measure intracellular glucose concentrations in living cells. We believe that this sensor can aid researchers interested in cellular metabolism.

## Resources

All TINGL plasmids can be obtained via Addgene (https://www.addgene.org/browse/article/28252279/)

## Data availability

All data and analysis scripts can be found online at 10.5281/zenodo.15174055.

## Author contributions

Conceptualization, D.B., J.G. and B.T.; Methodology, D.B.; Investigation, D.B. and A.T.; Writing, D.B., A.T., J.G. and B.T.; Resources, D.B.; Supervision, D.B.

## Supplementary information

**Figure S1.**
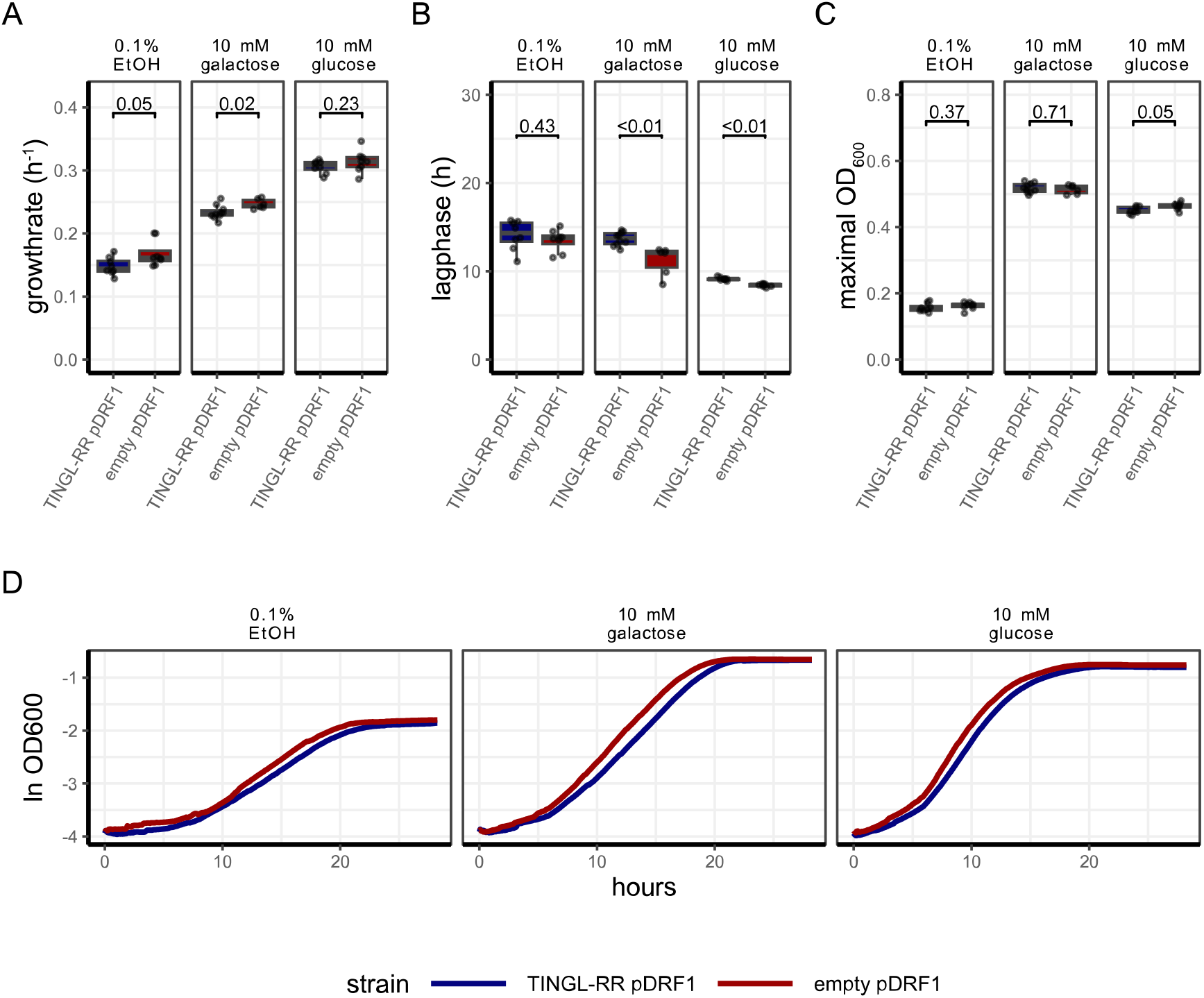
TINGL-RR can affect growth, although minimally. A) Growthrates of W303-1A expressing TINGL-RR in pDRF1 or the empty pDRF1 vector on 0.1% EtOH, 10 mM galactose and 10 mM glucose. B) lagphase (determined as the timepoint at which the cells obtain maximal growth) of W303-1A expressing TINGL-RR in pDRF1 or the empty pDRF1 vector on 0.1% EtOH, 10 mM galactose and 10 mM glucose. C) Maximal OD_600_ as a proxy of yield of W303-1A expressing TINGL-RR in pDRF1 or the empty pDRF1 vector on 0.1% EtOH, 10 mM galactose and 10 mM glucose. D) growth curves of W303-1A cells expressing TINGL-RR in pDRF1 or the empty pDRF1 vector. Lines show median optical density, colours indicate which plasmid is expressed. Boxplot indicates median with quartiles; whiskers indicate largest and smallest observations at 1.5 times the interquartile range. Dots show the technical replicates. P-values of a Wilcoxon test are shown between boxes.

**Figure S2.**
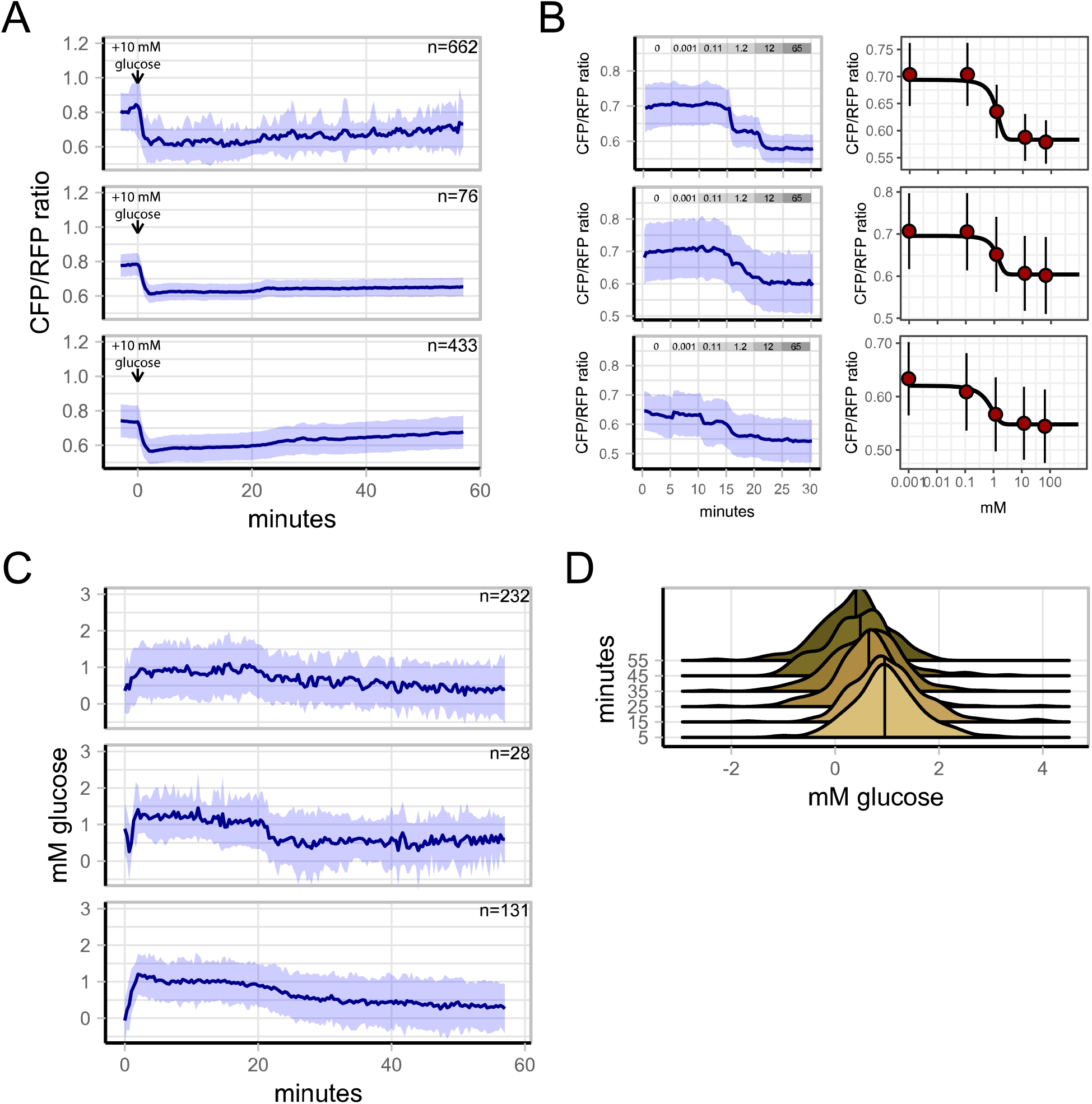
Ratios of TINGL-RR converted to actual intracellular glucose concentrations per biological replicate. A) CFP/RFP ratios of W303-1A cells grown on 1% EtOH to a 100 mM glucose pulse. Three biological replicates are shown. Lines show the mean CFP/RFP ratio, shades indicate SD. B) At the same day, an in-vivo dose-response was performed using the hexokinase mutant YSH757. Blue lines show mean CFP/RFP ratio, shades indicate SD. Black line shows Hillfit curve, dots and whiskers indicate mean CFP/RFP ratio with SD, respectively. C) The obtained dose-response curve for that day can be used to calculate the absolute glucose concentrations in cells. Note that the number of cells is different compared to panel A. This is because cells can have ratios outside the dose-response curve. This results in infinite values which cannot be depicted. The n values shown here are therefore the total number of cells, but at single timepoints the number of cells that show measurable CFP/RFP ratios can be lower. Lines show mean CFP/RFP ratio, shades indicate SD. D) distribution of glucose levels of all 3 replicates (shown in C) combined at various timepoints after the transitions (shown at the x-axis). Vertical line indicates the median of the distribution. Data obtained from three biological replicates.

